# The skull of *Tonatia bidens* Spix, 1823, found in ancient deposits of Abismo Ponta de Flecha Cave, Ribeira Valley, Southeastern Brazil

**DOI:** 10.1101/2025.08.11.669759

**Authors:** Artur Chahud

## Abstract

The Ribeira Valley, located in the southern part of São Paulo State, contains an extensive cave system where vertebrate fossils from the Late Pleistocene and Holocene have been found. The Abismo Ponta de Flecha Cave, a vertical cave rich in osteological material, is subdivided into lateral galleries known as *Jazidas*. Previous paleontological studies have described several extinct and extant species, but chiropterans were only briefly mentioned in the initial study of this cave, with only a single fragmented skull reported. This study presents that skull and briefly discusses the locality where it was found. The specimen belonged to an adult individual of *Tonatia bidens*, discovered in *Jazida* 6, an intermediate level located some distance from the cave entrance. Bat osteological remains are fragile and easily destroyed in high-energy environments; therefore, the preservation observed in this *Jazida* may indicate that the specimen died *in situ*, without being transported. Although this area experienced intense sediment remobilization in the past, *Jazida* 6 shows evidence of stability and minimal reworking activity, as indicated by the preservation of fragile specimens, sometimes still in their anatomical position at death.

## INTRODUCTION

The Ribeira Valley, in southern São Paulo State, is underlain by Precambrian carbonate rocks and hosts a complex system of caves. One of these, the Abismo Ponta de Flecha Cave, contains a vertebrate fossil assemblage from the Late Pleistocene and Holocene.

Despite the considerable paleobiological potential offered by the Ribeira Valley, due to its extensive system of limestone caves, the number of studies conducted in the area remains relatively small. The earliest paleontological research dates back to the early 20th century, beginning with AMEGHINO (1907), followed by studies in the 1970s and early 1980s by PAULA-COUTO (1973, 1979) and LINO et al. (1979), which primarily focused on documenting the local paleofauna. These works recorded fossils representative of the South American Pleistocene fauna preserved within caves, along with remains of numerous species still present in the modern fauna.

Only in the 21st century has a significant volume of paleontological research been carried out in the Ribeira Valley, with studies focusing on reptiles (CAMOLEZ & ZAHER, 2010), birds (CHAHUD, 2023b), mammals from both extant and extinct faunas (FERREIRA & KARMANN, 2002; CASTRO & LANGER, 2008, 2011; CHAHUD et al. 2023b; 2023c; 2024a; 2024b), and radiometric dating of fossil specimens (NEVES et al., 2007; HUBBE et al., 2013).

One of the most extensively investigated sites in the region is the Abismo Ponta de Flecha, a vertical cave subdivided into lateral galleries, locally referred to as *Jazidas*, each containing distinct sedimentary deposits and osteological material. The first comprehensive study was carried out by BARROS BARRETO et al. (1982), who investigated the geology, paleontology, and archaeology of the site. In addition to fossil remains, these authors reported archaeological artifacts suggesting a possible coexistence between extinct fauna and pre-colonial human populations.

Although the material recovered in the late 1970s was carefully numbered according to deposits, the site was only recently analyzed in detail from a paleontological perspective by CHAHUD (2021, 2022a; 2023a), CHAHUD et al. (2022, 2023a), and COSTA et al. (2025). Nevertheless, many specimens remain poorly studied. Among the least investigated mammalian groups are bats (Order Chiroptera).

Bats are the only mammals capable of active flight and represent the second most diverse order of the Class Mammalia, surpassed only by rodents in number of species. Although they are frequently recorded in caves and natural shelters, their skeletal remains are relatively rare in archaeological and paleontological sites. The work by BARROS BARRETO et al. (1982) reported the presence of four species: *Tonatia bidens* Spix 1823, *Carollia perspicillata* Linnaeus, 1758, *Pygoderma bilabiatum* Wagner, 1843, and *Sturnira lilium* Geoffroy, 1810, but no images of the specimens were published, and three of these species were not observed in subsequent studies.

Since these materials do not appear in cataloging records, it is possible that these specimens were not incorporated into the scientific collection of the Institute of Geosciences of the University of São Paulo in the early 1980s or that the material was misplaced.

Among the remaining osteological material, only a fragmented skull of *Tonatia bidens* was found. This specimen had been incorrectly identified (attributed to an indeterminate carnivore) and had never been the subject of a formal study. The aim of the present work is to describe this skull and briefly discuss the locality within Abismo Ponta de Flecha Cave where it was recovered.

## MATERIAL AND METHODS

The Abismo Ponta de Flecha Cave is a predominantly vertical cave located at the bottom of an ancient polygonal depression with centripetal drainage (KARMANN, 1994). This geological feature is hosted within Middle to Late Proterozoic carbonate rocks in southeastern Brazil (Figure 1), specifically in the municipality of Iporanga, southern São Paulo State.

**Figure 1.**
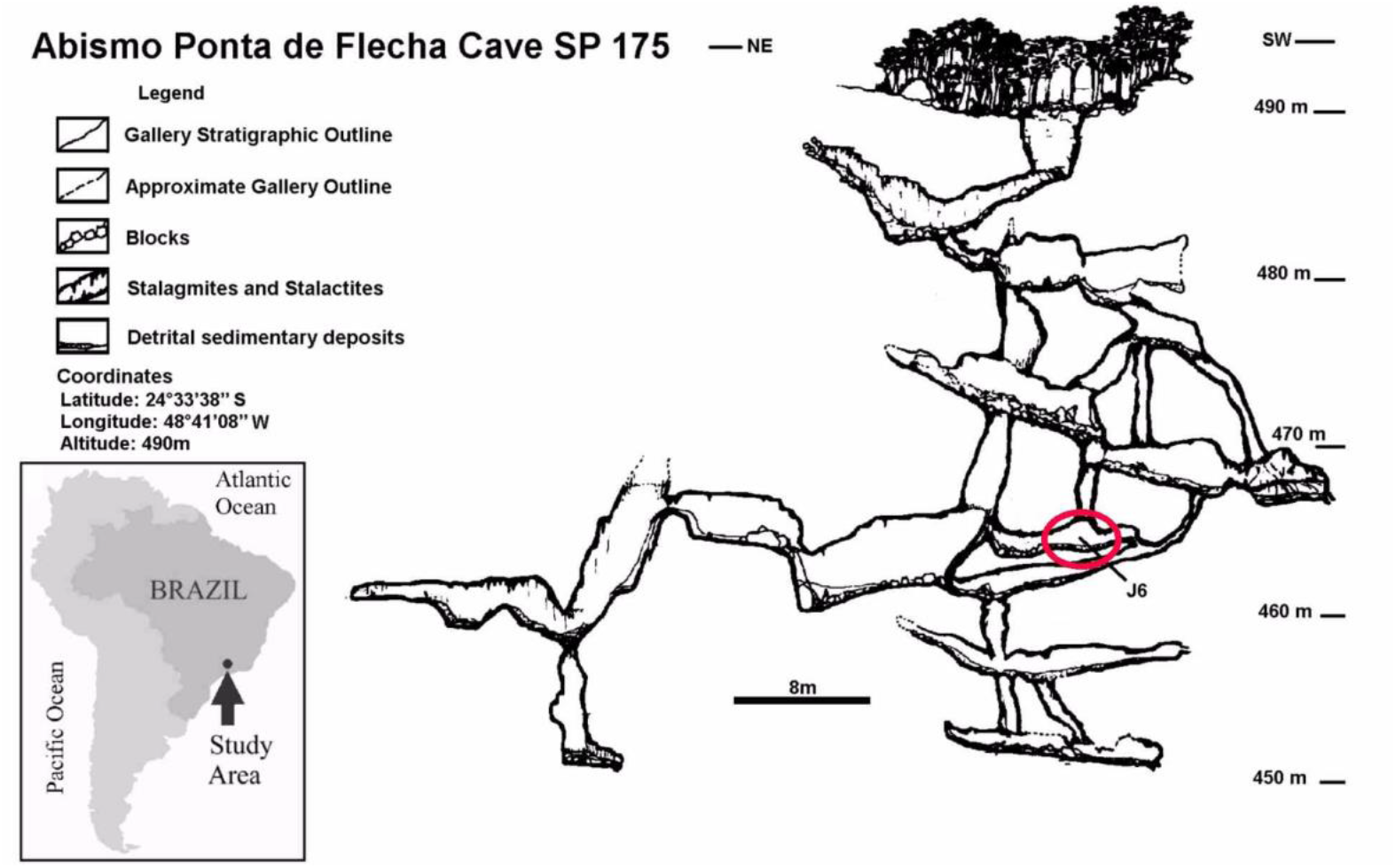
Schematic profile of Abismo Ponta de Flecha Cave, SP 175. Highlighting *Jazida* 6, where the specimen was found (Adapted from Barros-Barreto et al. 1982).

The analyzed material was collected between 1981 and 1982 and comprises over 1,400 samples of faunal and inorganic remains (BARROS-BARRETO et al., 1982). Excavation involved systematic removal of internal sediments, accompanied by documentation, transportation, and preliminary processing of the recovered material.

For excavation and recording purposes, the cave was divided into 11 *Jazidas*, corresponding to natural platforms. These *Jazidas* were numbered in descending order; however, this numbering does not necessarily reflect to the chronological sequence of excavation.

Each *Jazida* was further subdivided into concentrations and stratigraphic levels to systematize and enable precise spatial localization of the collected materials. Archaeological and paleontological techniques adapted to the cave environment were applied to ensure accurate recording and recovery (BARROS-BARRETO et al., 1982).

Specimen identification was conducted through comparison with reference collections and consultation of specialized literature. The specimen analyzed in this study is curated in the Systematic Paleontology Laboratory of the Department of Sedimentary and Environmental Geology at the Institute of Geosciences, University of São Paulo.

### CHARACTERISTICS OF THE ABISMO PONTA DE FLECHA CAVE

The development of the cave follows, for the most part, structural planes with a N30–40E trend and a 70–80SE dip, coinciding with the main structural direction of the region and with the valleys of the enclosing carbonate lens of the Ribeira Valley. The lithology is composed of two main units: calcitic metalimestones with marly layers, arranged concordantly within the core of a synformal structure composed of silty-clayey sediments with sandy intercalations (metasiltite and metarenite). Although originally located in a polygonal depression, the cavity is now situated at the top of a ridge (KARMANN, 1994).

Sedimentary deposition within the cave is highly irregular, both spatially and temporally. This variability is influenced by factors such as the location and morphology of the cavity, the position and shape of its entrance, the internal configuration, and the very nature of the accumulated sediments (BARROS-BARRETO et al. 1982).

As mentioned previously, the collectors divided the Abismo Ponta de Flecha Cave into 11 *Jazidas*. The term deposit, in this context, refers to a segment of the subterranean space that exhibits similar depositional processes, with concentrations of lithic material, osteological remains, or both, and may be of allochthonous or autochthonous origin. According to BARROS-BARRETO et al. (1982), the 11 *Jazidas* display distinct depositional characteristics. *Jazida* 6, where the specimen analyzed in this study was found, is composed of ancient sedimentary levels with collapsed blocks, indicating a phase of intense remobilization of the deposits.

### SISTEMATIC PALEONTOLOGY

Order: Chiroptera Blumenbach, 1779

Family: Phyllostomidae Gray, 1825

Genus: *Tonatia* Gray, 1827

*Tonatia bidens* Spix, 1823

Figure 2

**Figure 2.**
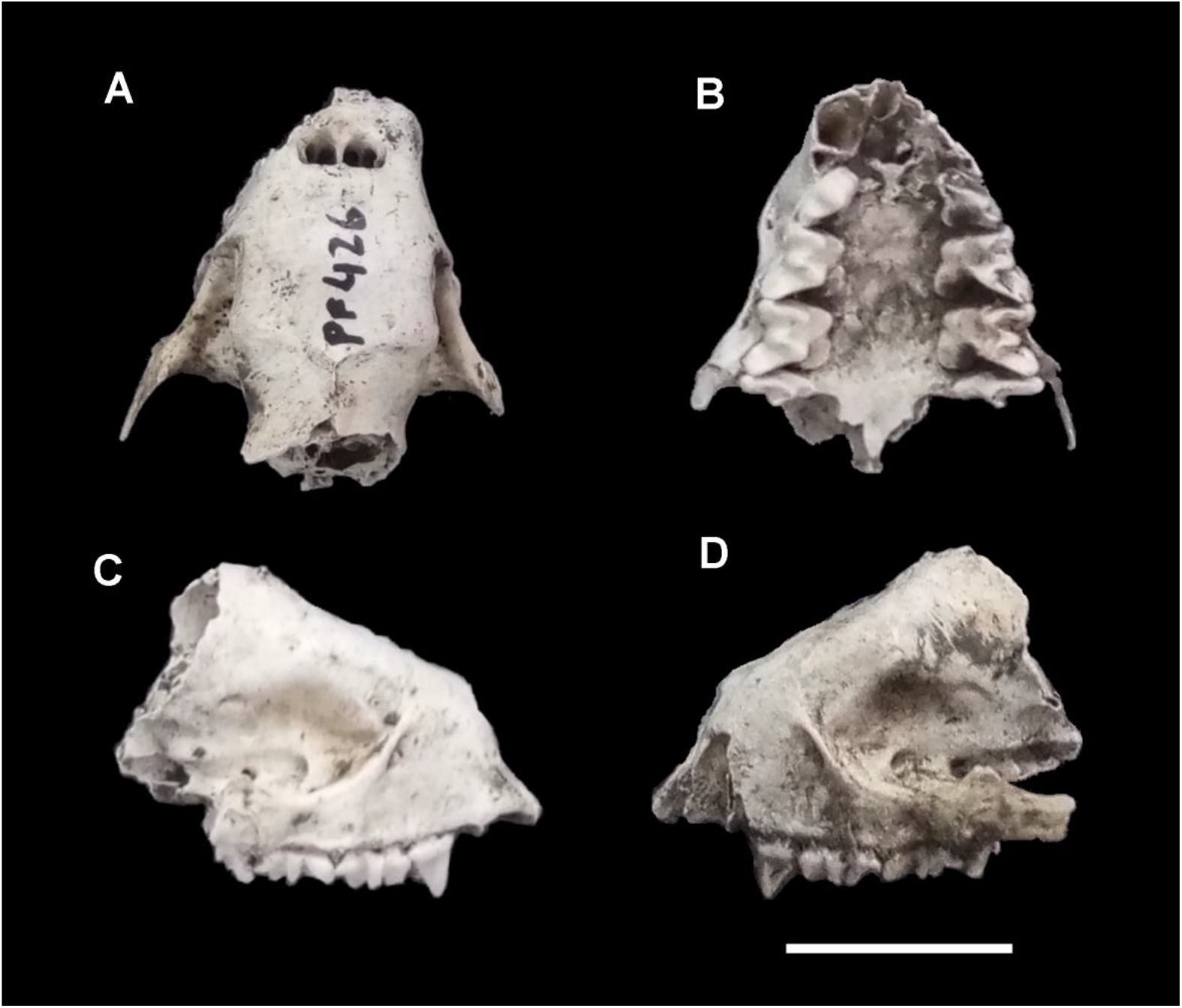
Fragmented skull of *Tonatia bidens* found in *Jazida* 6 of Abismo Ponta de Flecha Cave. A) Superior view; B) Occlusal view; C and D) Lateral views. Scale 10 mm.

### Material

Fragmented skull (GP/2C-174 – PF-426) recovered from *Jazida* 6 of Abismo Ponta de Flecha Cave (Figure 2).

### Distribution

*Tonatia bidens* has a geographic distribution ranging from Mexico to northern Argentina, including Paraguay and southeastern and eastern Brazil (BARQUEZ & DIAZ, 2016).

### Remarks

The specimen preserves only the anterior portion of the skull, including part of the nasal bones, the infraorbital margin, and the zygomatic arch (Figure 2A and 2B). Although it preserves a good part of the bony nasal opening, the anterior region of the skull lacks the prosthion, the incisors, canines, or the first premolar (Figures 2B). The right frontal portion is deformed and fragmented, possibly due to taphonomic processes.

The remaining dentition consists of the three upper molars on both sides, in addition to the second premolar (Figure 2C and 2D). The third molar is significantly smaller than the first two, and the second premolar has a greater relative height compared to the molars, a characteristic expected in members of the Family Phyllostomidae with an omnivorous or carnivorous diet and an elongated skull, such as the genera *Phyllostomus, Phylloderma*, and *Chrotopterus*.

Despite the fragmented skull, the dentition was key in identifying the specimen as *Tonatia bidens*, based on comparison with specimens from the Atlantic Forest, including specimens from the state of São Paulo (BRANDÃO & ZAHER, 2021; ROSA et al. 2022). The minimal wear on the permanent teeth indicates that the specimen was an adult, but not senescent.

## Discussion

*Tonatia bidens* is a solitary bat species that typically lives in small groups. It has an omnivorous diet and can consume a wide range of foods, including mosses (LUZ et al. 2024), fruits, insects (REDFORD & EISENBERG, 1992; ESBÉRARD & BERGALO, 2004; FELIX et al. 2013), birds, and small reptiles (CARVALHO et al. 2020). This dietary diversity reflects its ecological adaptability and trophic flexibility.

The genus *Tonatia* utilizes different types of shelter, range from natural caves to human-made constructions, and has been recorded in large urban areas (HADLER et al. 2018; ESBÉRARD & BERGALO, 2004; ROSA et al. 2022; GARBINO et al. 2022). It is quite common in the Atlantic Forest, and its occurrence in caves like Abismo Ponta de Flecha Cave is therefore not unexpected. However, it is surprising that the skull was found in such a deep area of the cave (Figure 1) and far from the main entrance. This fact raises the possibility that the individual became disoriented inside the cave or that the skull was transported to that location by internal water flows or biological agents (by carnivores or scavengers).

The presence of *Tonatia bidens* in Abismo Ponta de Flecha Cave had already been mentioned by BARROS BARRETO et al. (1982), but without any illustrations, detailed descriptions, or precise information of the provenance of the cited material. Therefore, it is not possible to confirm that the specimen analyzed here corresponds to one of those observed by those authors. It is also worth noting that in the collectors’ original record, the skull was incorrectly identified as belonging to a small carnivore, which may have contributed to its exclusion from later taxonomic studies.

### TAPHONOMY AND DEPOSITIONAL CHARACTERISTICS OF *JAZIDA* 6

As mentioned previously, *Jazida* 6 was described by BARROS BARRETO et al. (1982) as composed of ancient sedimentary platforms with collapsed blocks, indicating a phase of intense remobilization of the deposits. This description suggests that, at a certain point, the environment was subjected to high-energy conditions and intense reworking. However, the current depositional context in which the skull of *Tonatia bidens* was found likely does not correspond to this more dynamic phase.

Small vertebrates from Abismo Ponta de Flecha Cave were analyzed from a taphonomic perspective by CHAHUD (2022b), who, when studying the bone selection of marsupials, amphibians, and rodents, observed that the specimens were relatively recent. This is because intense reworking inside the cave tends to destroy more fragile bones, such as those of these small vertebrates. In the case of bats, whose bone structure is even more delicate, preservation is even less likely, which would explain the rarity of their remains at the site.

By definition, vertebrate skulls are among the last elements to be removed from a depositional environment and are considered part of residual deposits. This observation is exemplified by the Voorhies Groups (VOORHIES, 1969), in which Group I consists of small, light bones such as vertebrae, ribs, and phalanges; Group II is characterized by the predominance of long limb bones such as femurs, humeri, and tibias; and Group III includes mandibles and skulls, which tend to remain in their original location because they are less susceptible to transport.

I emphasize that the Voorhies Groups were proposed based on experiments with bones from medium-sized mammals, such as coyotes and sheep (VOORHIES, 1969). In the case analyzed at Abismo Ponta de Flecha Cave, the isolated presence of a small skull and a few bones of small animals suggests that transport events affected only small specimens, while larger animals remained in situ, as observed at *Jazida* 6.

*Jazida* 6, on the other hand, apparently differs from the other sites by providing the only specimen found in a death position at Abismo Ponta de Flecha Cave: an adult *Tayassu pecari*, whose skeleton was relatively well preserved (BARROS BARRETO et al. 1982). The specimen remained exposed for a long period without significant sedimentary cover, which can be inferred from the presence of encrustations on some bone elements.

The skull of *Tonatia bidens* was found in association with this *Tayassu pecari* specimen, along with other bones of small vertebrates, suggesting that, at the time of deposition, the *Jazida* 6 was already protected from high-energy events. This indicates that the conditions described by BARROS BARRETO et al. (1982) may no longer be active, or at least not with the same intensity observed in other locations in the cave, which allowed the preservation of more fragile materials in a relatively stable environment.

However, it is not possible to determine precisely when these lower-energy conditions began to prevail. The scarcity of osteological material and the absence of extinct species in *Jazida* 6 suggest that this environmental stabilization process is relatively recent.

Another hypothesis, which should be considered, is that the skull of *Tonatia bidens* and the associated small bones have a different origin than that of the *Tayassu pecari* specimen, possibly having been transported from other areas through earlier remobilization events. Alternatively, the *Tayassu pecari* may have reached *Jazida* 6 on its own, in a situation in which the animal entered the cave and was unable to exit, perishing there.

## CONCLUSIONS

The specimen recovered from Abismo Ponta de Flecha Cave belongs to an adult individual of the species *Tonatia bidens*. Represented by a fragmented skull, the specimen was located in an intermediate level of the cave (*Jazida* 6), situated a considerable distance from the entrance.

The presence of the specimen in this location may be associated with the individual’s disorientation during its incursion into the cave, ultimately leading to its death in situ. On the other hand, it is possible that the skull was transported from another area within the cave; however, this hypothesis is less likely, given the fragility of the material and the lack of evidence of long-distance or high-intensity transport.

*Jazida* 6 represents an area of Abismo Ponta de Flecha Cave where, during earlier phases of its formation, intense remobilization of sediment and internal elements occurred. However, taphonomic evidence indicates that this condition has not persisted into more recent times. Currently, this area appears to be one of the most stable sectors of the cave, as demonstrated by the presence of a relatively complete skeleton of *Tayassu pecari* in the anatomical position of death.

## Acknowledgements

The author thanks the Dr. Maria Mercedes Martinez Okumura, responsible for LEEH (Laboratory for Human Evolutionary Studies), Department of Genetics and Evolutionary Biology, Institute of Biosciences of the University of São Paulo for permitting the preparation of the fossils in her laboratory.

## Notes

### Competing Interest Statement

The authors have declared no competing interest.

